# Moving translational mass spectrometry imaging towards transparent and reproducible data analyses: A case study of an urothelial cancer cohort analyzed in the Galaxy framework

**DOI:** 10.1101/2021.08.09.455649

**Authors:** Melanie Christine Föll, Veronika Volkmann, Kathrin Enderle-Ammour, Konrad Wilhelm, Dan Guo, Olga Vitek, Peter Bronsert, Oliver Schilling

## Abstract

**Background:** Mass spectrometry imaging (MSI) derives spatial molecular distribution maps directly from clinical tissue specimens. This allows for spatial characterization of molecular compositions of different tissue types and tumor subtypes, which bears great potential for assisting pathologists with diagnostic decisions or personalized treatments. Unfortunately, progress in translational MSI is often hindered by insufficient quality control and lack of reproducible data analysis. Raw data and analysis scripts are rarely publicly shared. Here, we demonstrate the application of the Galaxy MSI tool set for the reproducible analysis of an urothelial carcinoma dataset.

**Methods:** Tryptic peptides were imaged in a cohort of 39 formalin-fixed, paraffin-embedded human urothelial cancer tissue cores with a MALDI-TOF/TOF device. The complete data analysis was performed in a fully transparent and reproducible manner on the European Galaxy Server. Annotations of tumor and stroma were performed by a pathologist and transferred to the MSI data to allow for supervised classifications of tumor vs. stroma tissue areas as well as for muscle-infiltrating and non-muscle invasive urothelial carcinomas. For putative peptide identifications, m/z features were matched to the MSiMass list.

**Results:** Rigorous quality control in combination with careful pre-processing enabled reduction of m/z shifts and intensity batch effects. High classification accuracy was found for both, tumor vs. stroma and muscle-infiltrating vs. non-muscle invasive tumors. Some of the most discriminative m/z features for each condition could be assigned a putative identity: Stromal tissue was characterized by collagen type I peptides and tumor tissue by histone and heat shock protein beta-1 peptides.

Intermediate filaments such as cytokeratins and vimentin were discriminative between the tumors with different muscle-infiltration status. To make the study fully reproducible and to advocate the criteria of FAIR (findability, accessibility, interoperability, and reusability) research data, we share the raw data, spectra annotations as well as all Galaxy histories and workflows. Data are available via ProteomeXchange with identifier PXD026459 and Galaxy results via https://github.com/foellmelanie/Bladder_MSI_Manuscript_Galaxy_links.

**Conclusion:** Here, we show that translational MSI data analysis in a fully transparent and reproducible manner is possible and we would like to encourage the community to join our efforts.

## BACKGROUND

Mass spectrometry imaging (MSI) is a label-free and untargeted method to generate spatial distribution maps for hundreds to thousands of molecules directly from a single tissue section. The most common MSI technique is based on matrix assisted laser desorption / ionization (MALDI) mass spectrometry and called MALDI MSI or MALDI imaging. It allows spatial resolution in the low micrometer range while preserving the integrity of the measured molecules such as proteins, peptides, metabolites and lipids. After the MALDI measurement, the tissue section remains amenable to histological staining, which can be compared to the measured molecular distributions. Molecular histology impacts many aspects of histopathological diagnostics and research and thus MSI is emerging as a powerful technology in translational studies [1,2]. In particular, the analysis of tumor tissues with pronounced cellular and morphological heterogeneity benefits from the spatially resolved MSI technology [3,4]. Common applications for MSI in cancer studies include tumor typing and subtyping [5–7], studying resection margins and tumor heterogeneity [8,9], and finding biomarkers for tumor diagnosis, prognosis or prediction [10–12].

The successes seen in translational MSI studies highlight the great potential for MSI in clinical settings, which require thorough quality control, good experimental design as well as standardized and reproducible experiments, analysis and reporting [2,13–16]. Despite their general importance for any omics-study, such aspects are only starting to become topics of research and developments in MSI. Recently, two studies emerged that demonstrated that standardized sample preparation protocols allow for reproducible MSI across several laboratories [14,15]. Suggestions for the inclusion of quality metrics into sample preparation protocol were made by Gustafsson et al. (use of internal peptide standards to measure and re-adjust mass accuracy [17]) and Erich et al. (implementation of quality controls for tryptic digestion efficiency [13]). In contrast, in most MSI studies the data analysis part is neither standardized, transparent nor reproducible, even though this part of an MSI experiment can be improved with the least effort. It requires publishing raw data and metadata as well as reporting the entire multistep analysis workflow with all fine grained parameters and settings in an accessible way.

We have previously established MSI tools in the Galaxy platform for reproducible MSI analyses [18]. Galaxy represents a highly suitable platform for reproducible biomedical data science allowing to track provenance, store tool names, versions and all set parameters for all analyses in publishable Galaxy history. Galaxy is accessible for every researcher, offers a graphical user interface, and comprises free access to thousands of pre-installed tools and large public computational resources. Galaxy also enables high levels of interoperability by implying tools of different omics domains which can be connected to build (multi-omics) workflows. Also, analysis histories and workflows can be shared privately, with collaboration partners or publicly, allowing full transparent and reproducible data analyses.

Here, we aim to showcase that a fully transparent and reproducible data analysis of a translational MSI cancer study is possible in the Galaxy framework. As a use case, we have imaged an urothelial tissue cohort comprising urothelial cancer, precursor lesions and benign tissues for their spatial tryptic peptide composition. Based on these tissues we established a classifier for two different tissue types, tumor and stroma. Considering the tumor, another classifier was built to distinguish between the two clinical relevant groups: muscle-infiltrating urothelial carcinoma and non-muscle invasive papillary urothelial carcinoma low grade (pTa low). The latter classifier could be applied to estimate the molecular risk of progression for three non-muscle invasive papillary urothelial carcinoma high-grade (pTa high) tissues. The classification of tumor areas from their surrounding stroma tissue is the key for tumor specific analysis. Currently, most MSI tumor studies rely on the manual annotation of tumor areas by a pathologist, which is a bottleneck in terms of available experts and time constraints, which could be overcome by applying automated classification of the two tissue types. The complete analysis including quality control, image co-registration, filtering regions of interest (ROIs), combining files, pre-processing, classification, and visualization was performed in a single platform: the European Galaxy server [19]. This allowed for the easy sharing of all analysis histories together with all raw and intermediate data to enable FAIR (findable, accessible, interoperable, and re-usable) data sharing and full transparency and reproducibility [20].

## METHODS

### Patient cohort

49 bladder tissue specimens from 47 patients were collected during transurethral resection at the University Medical Center in Freiburg. The study was approved by the Ethics Committee of the University Medical Center Freiburg (no. 491/16). All patients gave written informed consent. Before study inclusion, all patient data were pseudonymized.

Bladder tissue specimens were formalin-fixed directly after surgical removal and paraffin-embedded as described previously [21]. All tissue specimens were reviewed by two experienced pathologists. Biopsie tissue cores with 2 mm diameter were extracted from each formalin-fixed paraffin embedded (FFPE) tissue block and randomly assembled into two FFPE tissue microarrays (TMA) blocks. The following tissue cores were included into the TMA: Muscle-infiltrating urothelial cancer (n=12), non-muscle invasive papillary urothelial carcinoma high-(pTa high, n=5) / low grade (pTa low, n=20), carcinoma in situ (pTis, n=2), and papillary urothelial neoplasm of low malignant potential (PUNLUMP, n=2), as well as inflammatory bladder specimens (n=8). 6 µm thick sections were sliced with a microtome and mounted onto indium tin oxide (ITO) coated glass slides (Bruker Daltonik, Bremen, Germany).

### MSI sample preparation

Tissue deparaffinization was performed in xylol and ethanol/water solutions as described previously [21]. Tissue sections were rinsed twice in 10 mM ammonium bicarbonate (NH4HCO3) for 1 min. Antigen retrieval was performed in citric acid monohydrate pH 6.0, in a steamer for 1 h at approximately 100 °C [22]. Rinsing in ammonium bicarbonate was repeated twice and the samples were air dried afterwards. TPCK treated Trypsin (Worthington, Lakewood, NJ, USA) was sprayed onto the tissue sections using iMatrixSpray (Tardo Gmbh, Subingen, Switzerland); 60 mm height, 1 mm line distance, 180 mm/s speed, 0.5 µl/cm3 density, 10 cycles, 10 s delay [23]. Digestion was performed for 2 h at 50°C in a digestion chamber with 97% humidity maintained by a saturated potassium sulfate (K2SO4) solution [14]. 10 mg/ml alpha-cyano-4-hydroxycinnamic acid (CHCA, Sigma-Aldrich, Munich, Germany) matrix was prepared in 50% (v/v) acetonitrile and 1% (v/v) trifluoroacetic acid. Matrix solution was mixed 12:1 (v/v) with an internal calibrant mix containing 0.08 µg/ml Angiotensin I (Anaspec, Seraing, Belgium), 0.04 µg/ml Substance P (Anaspec, Seraing, Belgium), 0.15 µg/µl [Glu]-Fibrinopeptide B (Sigma-Aldrich, Munich, Germany), and 0.30 µg/µl ACTH fragment (18-39) (Abcam, Cambridge, UK) [17]. The matrix-calibrant mixture was sprayed onto the tissue sections using iMatrixSpray; 60 mm height, 1 mm line distance, 180 mm/s speed, 0.5 µl/cm3 density, 20 cycles, 5 s delay.

### MSI data acquisition

Tissue sections were measured with a 4800 MALDI-TOF/TOF Analyzer (Applied Biosystems, Waltham, MA, USA) using the 4000 Series Explorer software (Novartis and Applied Biosystems) to set instrument parameters. A squared region was imaged with 150 µm raster step size, averaging 500 laser shots per spectrum in a mass range from 800 to 2300 m/z in positive ion reflectron mode. Before starting the imaging measurement, internal calibrants in a spectrum outside the tissue region were used for m/z re-calibration.

### Tissue staining and annotation

Matrix was removed from the slides by rinsing with 70 % ethanol after MSI measurement. Afterwards, hemalum staining of the measured tissue was performed by immersing the tissue sections in Mayer’s acid Hemalum solution (Waldeck, Münster, Germany) for 1 minute and rinsing with water for 1 minute. Dehydration was performed with four short incubations in 100% ethanol and 2 incubations in xylol. Stained tissues were scanned at x20 magnification. A pathologist (KEA) annotated a coherent area within the largest tumor and stroma regions in photoshop CS5 (Adobe, San Jose, USA). Only annotated spectra were considered for further analysis.

### MSI quality control, data handling and pre-processing

Analyze7.5 files were uploaded to the European Galaxy server [24], where the complete analysis was performed and afterwards published [18,19]. First, a quality control with the MSI qualitycontrol tool (m/z of interest: four internal calibrants, ppm range: 200) was performed to ensure sufficient quality of the data and to find appropriate parameters for the following pre-processing steps. A previously published Galaxy workflow [18] was slightly modified and applied for co-registration of the stained image and the MSI image for each TMA separately. Six visually determined characteristic tissue spots were used as teachmarks for affine transformation. The obtained warping matrix was applied to extract the coordinates corresponding to the annotated regions from the MSI data leading to 2169 tumor and stroma specific spectra, while all pTis and PUNLUMP spectra were removed. Both files were binned in 50 ppm m/z steps and cut to their common m/z range 800 to 2300 in the ‘MSI preprocessing’ tool (method: m/z binning, width of bin: 50, unit for bin: ppm, select m/z options: change m/z range, minimum value for m/z: 800, maximum value for m/z: 2300 and combined into one dataset using the ‘MSI combine’ tool (Optional annotation of pixels with tabular files: TMA1 annotations, TMA2 annotations). The Cardinal (v 2.6.0) [25] based ‘MSI preprocessing’ tool was used for pre-processing: gaussian smoothing (window: 8, standard deviation: 2), baseline reduction (blocks: 750), m/z alignment (tolerance: 200 ppm), peak picking (signal to noise: 5, blocks: 600, window: 10), alignment (tolerance: 200 ppm) and filtering (frequency: 0.01) to obtain a common m/z peak list. The m/z peak list was used to extract the original peptide intensity from the smoothed and baseline reduced dataset by peak binning (tolerance: 200 ppm) in the ‘MSI preprocessing’ tool. Mass re-calibration (tolerance: 200 ppm) was performed based on the three internal calibrants within the m/z range and the most abundant tryptic autolysis peptide (m/z 405.42) using the align spectra function of the MALDIquant peak detection tool (tolerance: 0.0002, don’ t throw an error when less than 2 reference m/z were found in a spectrum: Yes, If TRUE the intensity values of MassSpectrum or MassPeaks objects with missing (NA) warping functions are set to zero: Yes, Should empty spectra be removed: Yes). Afterwards the processed imzML data was converted into a continuous file with the ‘MSI preprocessing’ tool (Processed imzML file: Yes, mass accuracy to which the m/z values will be binned: 0.005, unit of the mass accuracy: mz; preprocessing method: peak filtering, minimum frequency 0.01). Potential contaminant m/z features were removed with the ‘MSI filtering’ tool (Select m/z feature filtering option: remove m/z, tabular file with m/z features to remove: potential contaminant list, window in which all m/z will be removed: 200, units: ppm). The potential contaminant list was built based on the internal calibrants as well as CHCA matrix peaks and bovine trypsin peptides. The m/z of the latter two were obtained from the MALDI contaminant list published by Keller [26]. Finally, intensity normalization to the total ion current (TIC) of each spectrum was performed in the ‘MSI preprocessing’ tool. Between and after the pre-processing steps eight times a quality control was performed with the ‘MSI qualitycontrol’ tool using the three internal calibrants, a 200 ppm range and spectra annotation information to summarize either the properties of each TMA or of each patient tissue core.

### MSI statistical modelling, visualizations and identification

The pre-processed file was subjected to spectra classification using Cardinal’s spatial shrunken centroids (SSC) algorithm [27] in the ‘MSI classification’ tool. For tumor vs. stroma classification, stroma of non-malignant tissues and tumor tissues were not separated. All 39 patients were split randomly 80:20 into training (n=31) and test group (n=8). The patients of the training group were split into ten random groups. The scikit learn [28] based Split Dataset tool was used for all the patient grouping steps and the ‘MSI filtering’ tool in order to separate all training and test spectra into separate imzML files. First, 10-fold cross validation was performed on the training file in the ‘MSI classification’ tool (Pixel coordinates and their classes: file from Split Dataset tool that contains the spectra conditions and folds of the training data, select the method for classification: spatial shrunken centroids, analysis step to perform: cvApply, write out best r and s values: yes, r: 2, s: 0,2,4,6,8,10,12,14,16,18,20,22,24,26,28,30,32,34,36,38,40, method to use to calculate the spatial smoothing kernels: adaptive) to find optimal classification parameters. The optimized parameters (r=2, s=18) were applied to build a classifier on the training data with the ‘MSI classification’ tool (Pixel coordinates and their classes: file from Split Dataset tool that contains the spectra conditions of the training data, select the method for classification: spatial shrunken centroids, analysis step to perform: spatial shrunken centroids, r: 2, s: 18, method to use to calculate the spatial smoothing kernels: adaptive, Results as .RData output: yes). The classifier obtained as .RData file was then applied to the test data in the ‘MSI classification’ tool (Analysis step to perform: prediction, which classification method was used: SSC_classifier, load annotations: use annotations, load tabular file with pixel coordinates and the new response: file from Split Dataset tool that contains the spectra conditions of the test data).

For muscle-infiltrating vs. non-muscle invasive pTa low-grade tumor classification, only tumor ROIs from muscle-infiltrating urothelial cancer and non-muscle invasive low-grade papillary urothelial cancer were included into the analysis (more details in table1). Patients were randomly assigned 80:20 into training (n=20) and test group (n=6). The training group was further split into five random groups and 5-fold cross validation was performed to find optimal classification parameters. Again, the scikit learn based Split Dataset tool was used for all the patient grouping steps and the ‘MSI filtering’ tool in order to separate all training and test spectra into separate imzML files. First, 5-fold cross validation was performed on the training file in the ‘MSI classification’ tool (Pixel coordinates and their classes: file from Split Dataset tool that contains the spectra conditions and folds of the training data, select the method for classification: spatial shrunken centroids, analysis step to perform: cvApply, write out best r and s values: yes, r: 2, s: 0,2,4,6,8,10,12,14,16,18,20, method to use to calculate the spatial smoothing kernels: adaptive) to find optimal classification parameters. The optimized parameters (r=2, s=4) were used to build a classifier on the training data with the ‘MSI classification’ tool (Pixel coordinates and their classes: file from Split Dataset tool that contains the spectra conditions of the training data, select the method for classification: spatial shrunken centroids, analysis step to perform: spatial shrunken centroids, r:2, s:4, method to use to calculate the spatial smoothing kernels: adaptive, Results as .RData output: yes). The classifier obtained as .RData file was then applied to the test data in the ‘MSI classification’ tool (Analysis step to perform: prediction, which classification method was used: SSC_classifier, load annotations: use annotations, load tabular file with pixel coordinates and the new response: file from Split Dataset tool that contains the spectra conditions of the test data).

Furthermore, this classifier was applied to the tumor ROIs of the three non-muscle invasive high-grade papillary urothelial cancers to predict their invasiveness potential by using the ‘MSI classification’ tool (Analysis step to perform: prediction, which classification method was used: SSC_classifier, load annotations: use annotations, load tabular file with pixel coordinates and the new response: file from Split Dataset tool that contains the spectra conditions of the high-grade tumors). The most discriminative m/z features were selected according to the highest t-statistic values and their abundances in the different groups were visualized. Ion images were plotted with the ‘MSI mz images’ tool (plusminus m/z: 0.25, contrast enhancement ‘histogram’) on the binned, filtered, and combined data, which was TIC normalized in the ‘MSI preprocessing’ tool in a separate step only for visualization purposes. Average mass spectra plots per group were generated from binned, filtered, combined and smoothed MSI data with the ‘MSI plot spectra’ tool (choose spectra: plot single spectra, load tabular file with pixel coordinates: combined spectra annotations, separate plot per spectrum or overlaid plot with average spectra per annotation group: overlaid spectra plots, zoomed in m/z range: tabular file with mz of interest, m/z value to subtract from m/z values in tabular file: 1, m/z value to add to m/z values in tabular file: 4, load tabular file with m/z values: file with top mz value per condition). All m/z features that were part of one of the two classifiers (t-statistic value above zero) were matched with the Join two files tool (Choose the metrics of your distance: ppm, allowed distance between the two values that will trigger a merge: 200) to the downloaded MSiMass list [29] to obtain putative identifications. For figures 2 to 5, pdf files from Galaxy were imported into Adobe Illustrator CS2 to arrange subfigures and adjust the label sizes.

## RESULTS

### Overview of the urothelial cancer cohort

The urothelial cancer cohort consisted of two TMAs comprising 49 bladder tissue cores derived from 47 patients. Two tissue cores were lost during sample preparation and in three tissue cores, neither tumor nor stroma regions were withdrawn during TMA construction. Due to the insufficient sample size number one pTis and two PUNLUMP were excluded from the analysis. The exclusion of these tissues led to a final cohort of 39 tissue cores from 39 patients (Table 1) and 2169 mass spectra out of which 1076 were annotated as tumor and 1093 as stroma.

**Table 1:**
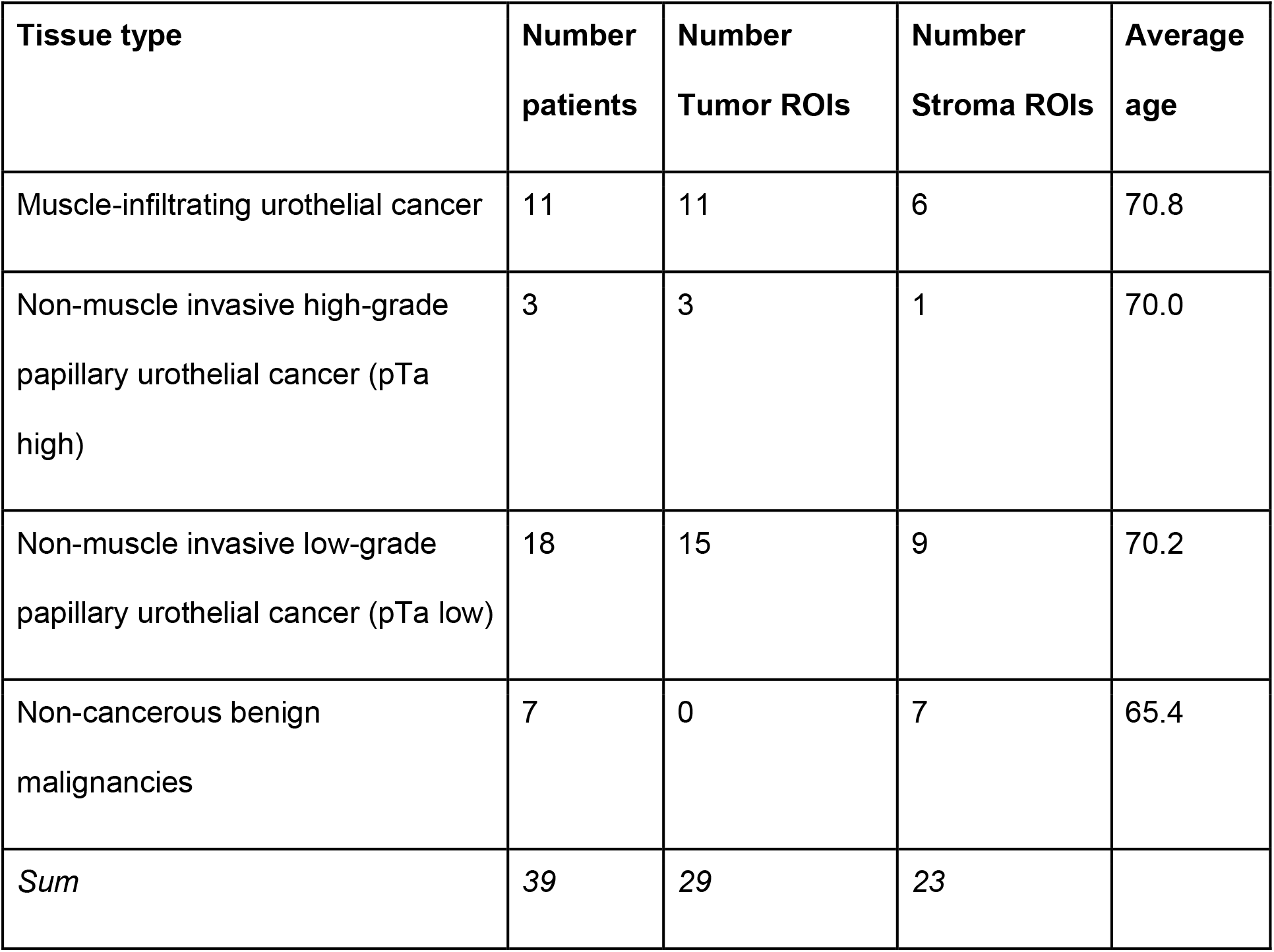
Overview of the patients and regions of interest (ROIs) of the urothelial cancer cohort

| **Tissue type** | **Number patients** | **Number Tumor ROIs** | **Number Stroma ROIs** | **Average age** |
| --- | --- | --- | --- | --- |
| Muscle-infiltrating urothelial cancer | 11 | 11 | 6 | 70.8 |
| Non-muscle invasive high-grade papillary urothelial cancer (pTa high) | 3 | 3 | 1 | 70.0 |
| Non-muscle invasive low-grade papillary urothelial cancer (pTa low) | 18 | 15 | 9 | 70.2 |
| Non-cancerous benign malignancies | 7 | 0 | 7 | 65.4 |
| *Sum* | *39* | *29* | *23* |  |

### Transparency and reproducibility of the MSI data analysis in the Galaxy framework

Both TMAs were imaged for tryptic peptides, hematoxylin and eosin stained and annotated for tumor and stroma ROIs. Raw data and spectra annotation information have been published via the PRIDE repository (identifier:PXD026459) [30]. The complete data analysis was performed on the European Galaxy server and was separated into seven different analysis histories, to keep the histories clearly arranged according to the different analysis steps: Co-registrations, data preparation and preprocessing, classifications, visualizations, and identification (Fig. 1a). To achieve full reproducibility and transparency of the study we published all Galaxy histories, which contain raw and intermediate files together with the tool name, tool version and all set parameters. For each of the first five analysis steps, Galaxy workflows were built and published to enable re-running the same analysis in a standardized and automated way. The pre-processing workflow is depicted as an example in Fig. 1b.

**Fig. 1:**
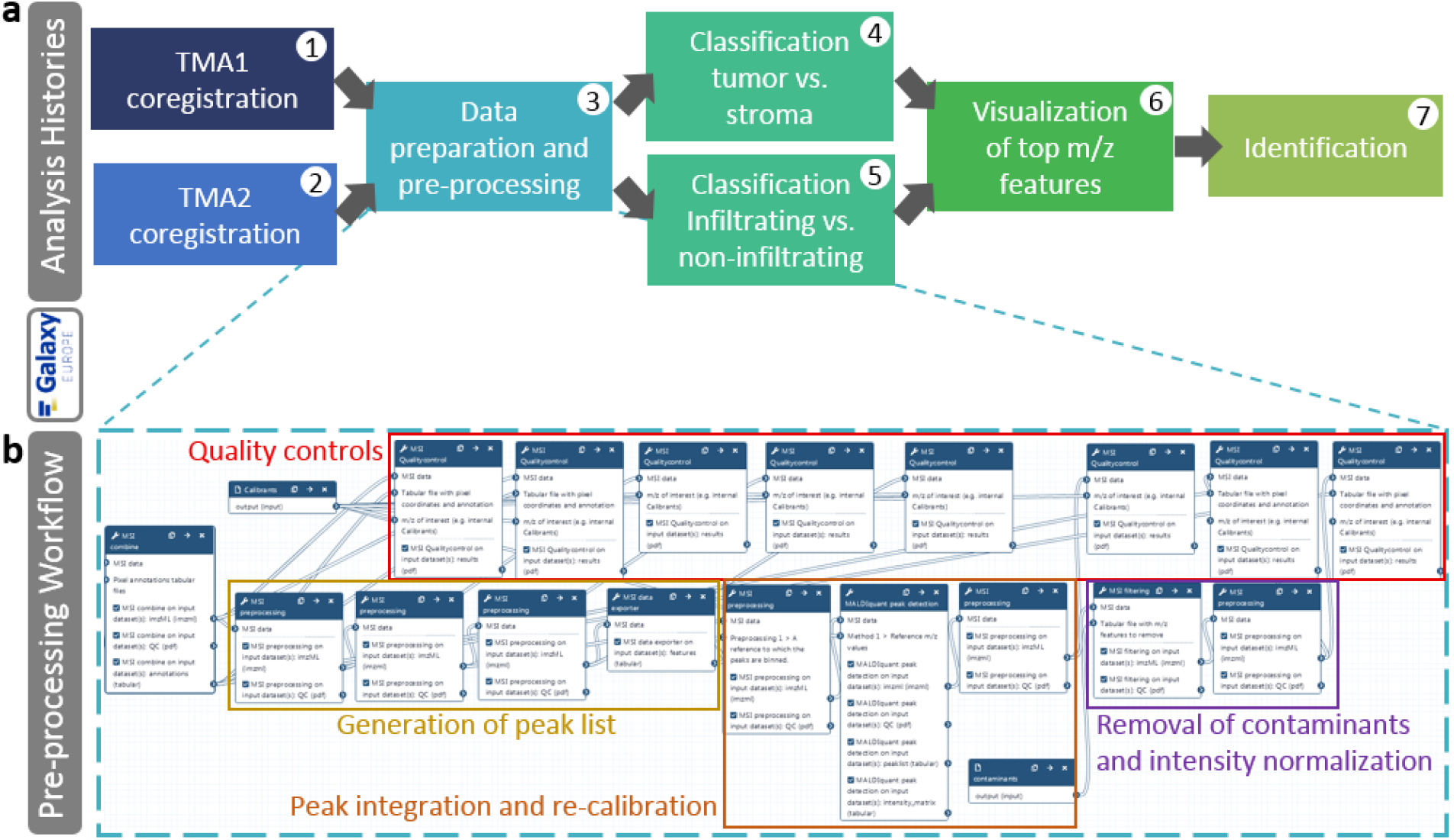
Overview of the data analysis pipeline. a) Overview of the performed analysis steps and their corresponding Galaxy histories. b) Galaxy workflow for pre-processing was built in a stepwise manner and combined with regular quality control steps. All Galaxy histories and workflows are published, links to them can be found in the ‘Availability of data and materials’ section.

### Quality control and preprocessing

The acquired data showed pronounced intensity batch effects and m/z shifts, which could be removed through careful adjustment of the preprocessing steps. Key to observe and overcome these technical issues was the usage of internal calibrants [17] together with the Galaxy ‘MSI qualitycontrol’ tool, which generated more than 30 different descriptive plots. Both TMAs showed systematically increasing m/z values for the internal calibrants during the course of the measurement (Fig. 2a). This suggests that the TOF tube of the near-antique mass spectrometer, which was not built to acquire tens of thousands of spectra in a row, heated up during the measurement. These m/z shifts could be removed by aligning each spectrum to the mean spectrum and re-calibrating the m/z positions via the internal calibrants (Fig. 2b).

**Fig. 2:**
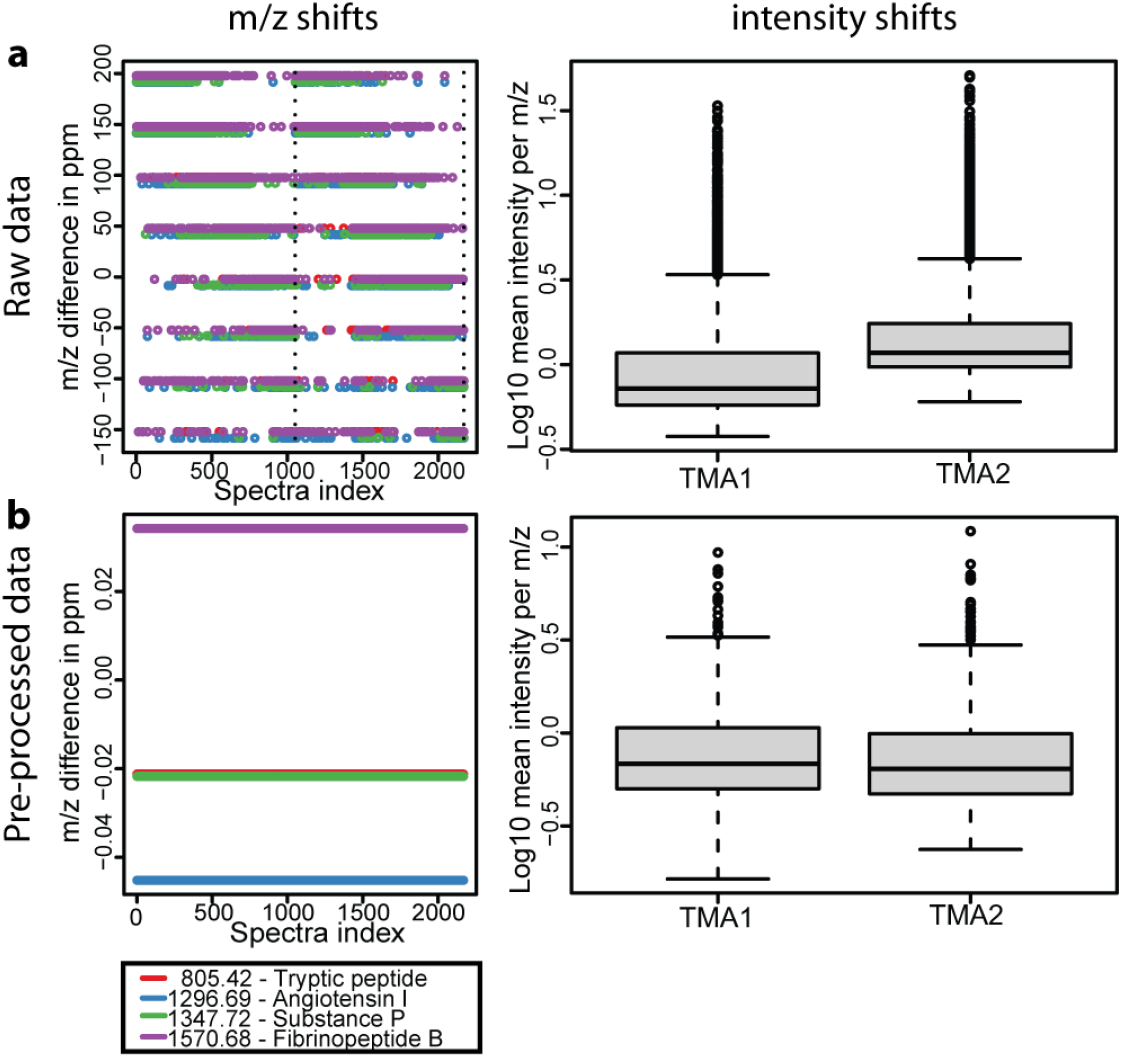
Data properties accessed by the ‘MSI qualitycontrol’ tool. a) Mass and intensity shifts before pre-processing. b) Mass and intensity shifts could be reduced through careful adjustment of pre-processing steps and parameters.

Intensity batch effects were observed between the two measurements with higher intensities in TMA2 (Fig. 2a). As the baseline was already removed during data acquisition, TIC normalization could not be performed on the raw data as suggested by Deininger et al. [31]. Instead we were able to reduce the batch effects (Fig. 2b) by performing TIC normalization after peak picking and contaminant removal as suggested by Fonville [32].

### Classification of Tumor and Stroma Spectra

In several cancers, tumor cells are intermingled or surrounded by connective tissue, the so-called tumor stroma, which is part of the tumor microenvironment. To distinguish tumor and stroma tissue types, we have built a classifier, which reached 94 % on the training and 99 % on the test datasets with high sensitivity and specificity (Table 2). To avoid overfitting, we generated training and test datasets by splitting patients randomly into the two groups and thus guarantee that all spectra of the same patient are present only in one of the two groups. Despite these precautions, the classification accuracy is likely too optimistic due to the small amount of samples. During classification, feature selection was performed by shrinking the number of m/z features that are included into the classifier to a minimum. 77 m/z features were included in the classifier (t-statistic > 0) out of which 37 were describing tumor and 40 stroma spectra (Fig. 3a). m/z 901.49 and 868.47 had the highest t-statistics for tumor and stroma respectively and were therefore the most discriminative m/z features (Fig.3b, c).

**Table 2:** Classification results tumor vs. stroma tissues

|  | Accuracy | Sensitivity | Specificity |
| --- | --- | --- | --- |
| Training | 0.94 | 0.94 | 0.94 |
| Test | 0.99 | 1 | 0.98 |

**Fig. 3:**
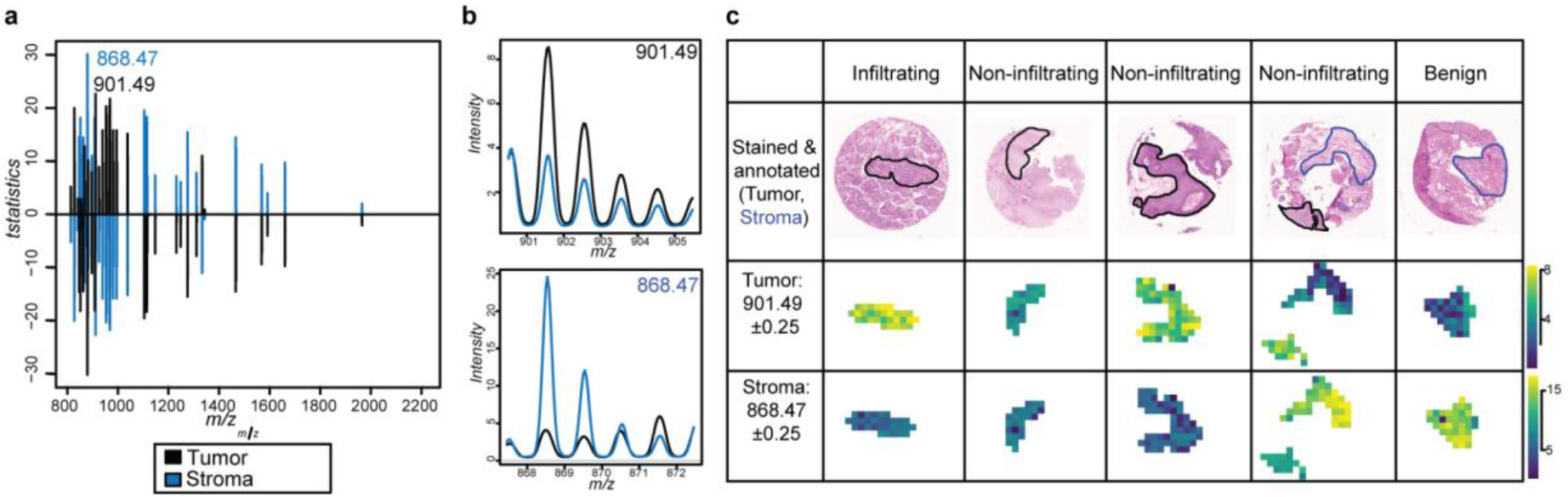
Classification results tumor vs. stroma and visualizations of the top m/z features of tumor and stroma respectively. a) Classification included feature selection based on t-statistics values above zero reveals 37 tumor specific and 40 stroma specific m/z features. b) Average mass spectra plots for the top m/z feature per group on binned, filtered, combined and smoothed MSI data. c) Ion images of the top m/z feature per group were plotted on five tissue cores with contrast enhancement ‘histogram’ on binned, filtered, combined and TIC normalized data.

### Classification of infiltration behavior

Next, we were interested in classifying tumors according to their infiltration status. Only spectra corresponding to muscle-infiltrating urothelial cancer (n = 11, 731 spectra) and non-muscle invasive low-grade papillary urothelial cancer (n = 15, 312 spectra) were included to compare both tumor subtypes. Classification accuracies for the training data were 96 % and 99 % for the test data with high sensitivity and specificity (Table 3). Again, the classification accuracy is likely too optimistic due to the small amount of samples. The classifier included 35 m/z features to classify muscle-infiltrating and 36 m/z features to classify non-muscle invasive tumors (Fig. 4a). The m/z feature 944.53 was the most discriminative for muscle-infiltrating tumor and m/z 1104.57 for non-infiltrating tumors (Fig. 4b, c).

**Table 3:**
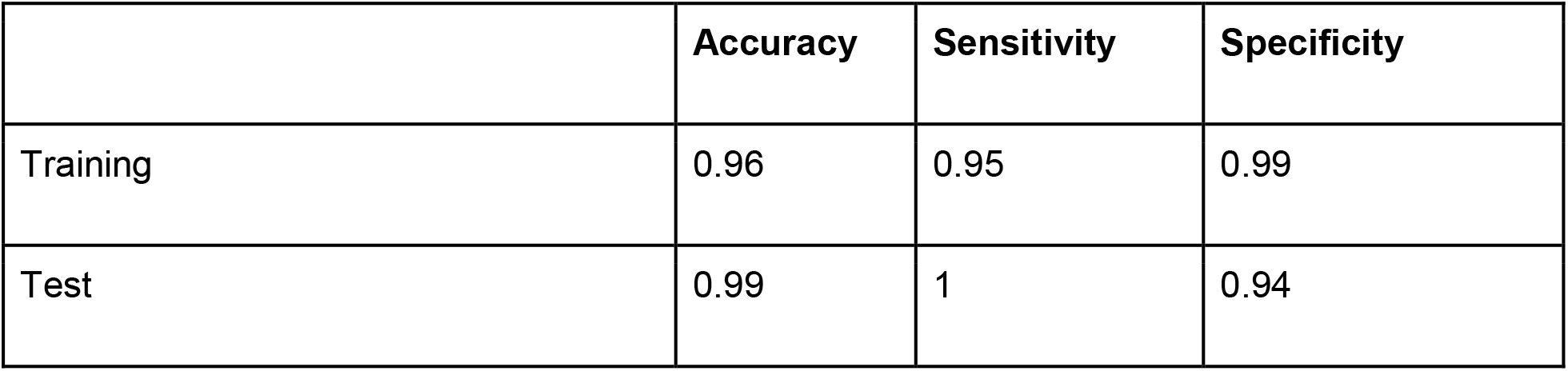
Classification results muscle-infiltrating vs. non-infiltrating carcinomas

**Fig. 4:**
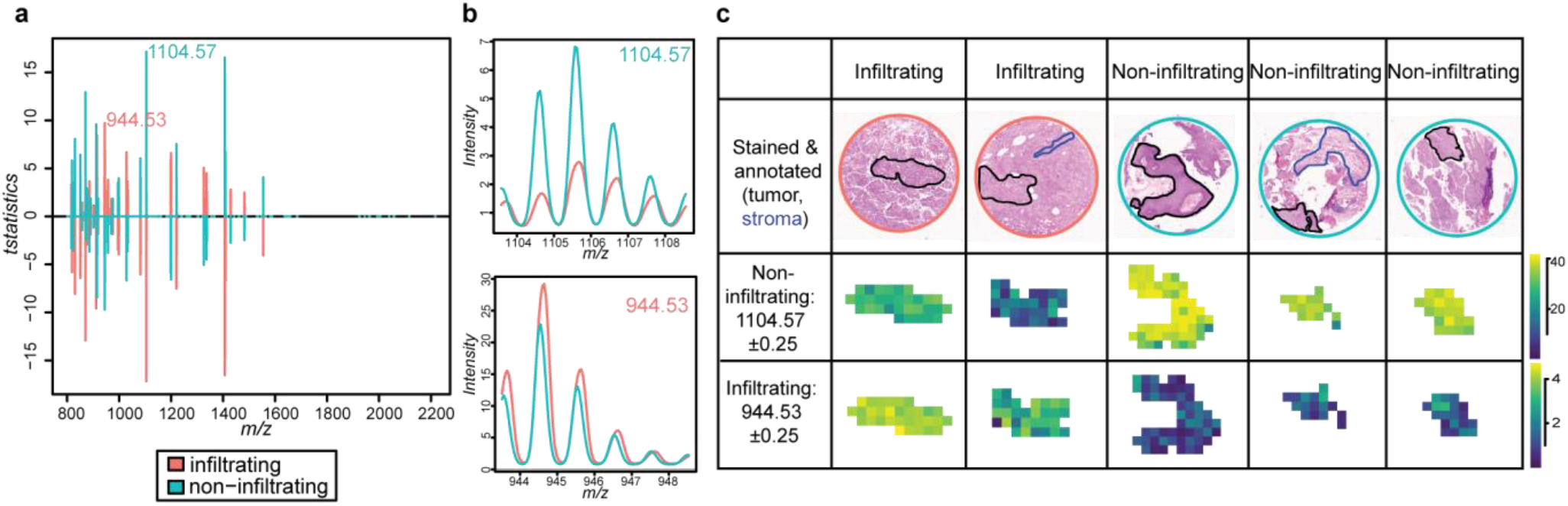
Classification results muscle-infiltrating vs. non-muscle infiltrating tumors and visualizations of their top m/z features. a) Classification included feature selection based on t-statistics values above zero reveals 35 muscle-infiltrating specific and 36 non-muscle infiltrating specific m/z features. b) Average mass spectra plots for the top m/z feature per group on binned, filtered, combined and smoothed MSI data. c) Ion images of the top m/z feature per group were plotted on five tissue cores with contrast enhancement ‘histogram’ on binned, filtered, combined and TIC normalized data.

### Prediction of muscle-infiltration potential of high-grade carcinomas

pTa high-grade urothelial cancers are not muscle invasive but are considered high-risk tumors as their risk of progression ranges from 15% to 40% and is thus much higher compared to pTa low-grade cancers [33]. The tissue cohort included three non-muscle invasive high-grade papillary urothelial cancer tissues (33 spectra), which were not included into the classification analysis because of their low sample number and being present only in one of the two TMAs.

Instead, we determined their muscle-infiltration potential by classifying them with the previously built classifier into muscle-infiltrating and non-muscle invasive cancers. With a total of only three patients, this analysis step is rather illustrative. The majority of spectra of all three tissue cores was classified as non-muscle invasive but in one tissue 2 out of 15 spectra were classified as muscle-infiltrating and several other spectra were classified only with low probabilities as non-muscle invasive, suggesting that this cancer might have the molecular potential to transition into a muscle-infiltrating cancer (Fig. 5). Unfortunately this hypothesis could not be verified by clinical data because the patient was lost to follow up.

**Fig. 5:**
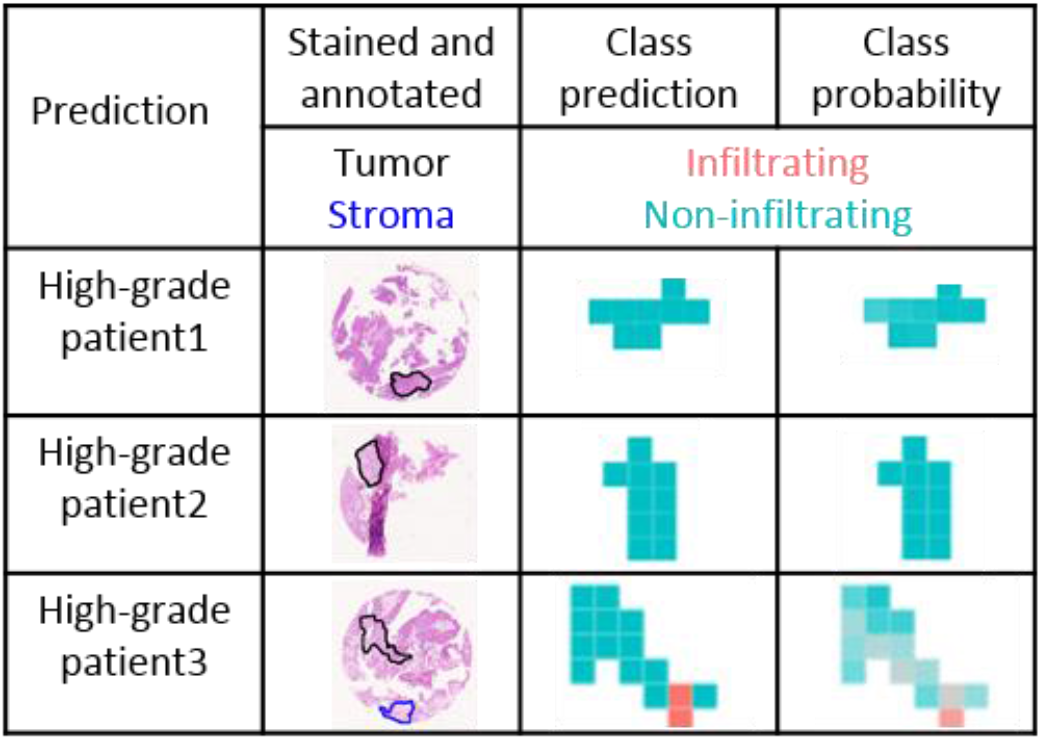
Prediction of high-grade urothelial carcinomas as non-muscle infiltrating and muscle-infiltrating cancers.

### Assigning identities to m/z features

To obtain an idea about the identity of the measured peptides, we assigned tentative identifications via the MSiMass list [29]. Out of 123 unique m/z features that were part of the two classifiers (t-statistics >0), 16 were matched to an entry of the MSiMass list within 200 ppm mass tolerance (Table 4). Most tentative collagen peptides were found in stromal regions and most keratin peptides in tumor regions, which is their expected location in tumor tissues.

**Table 4:**
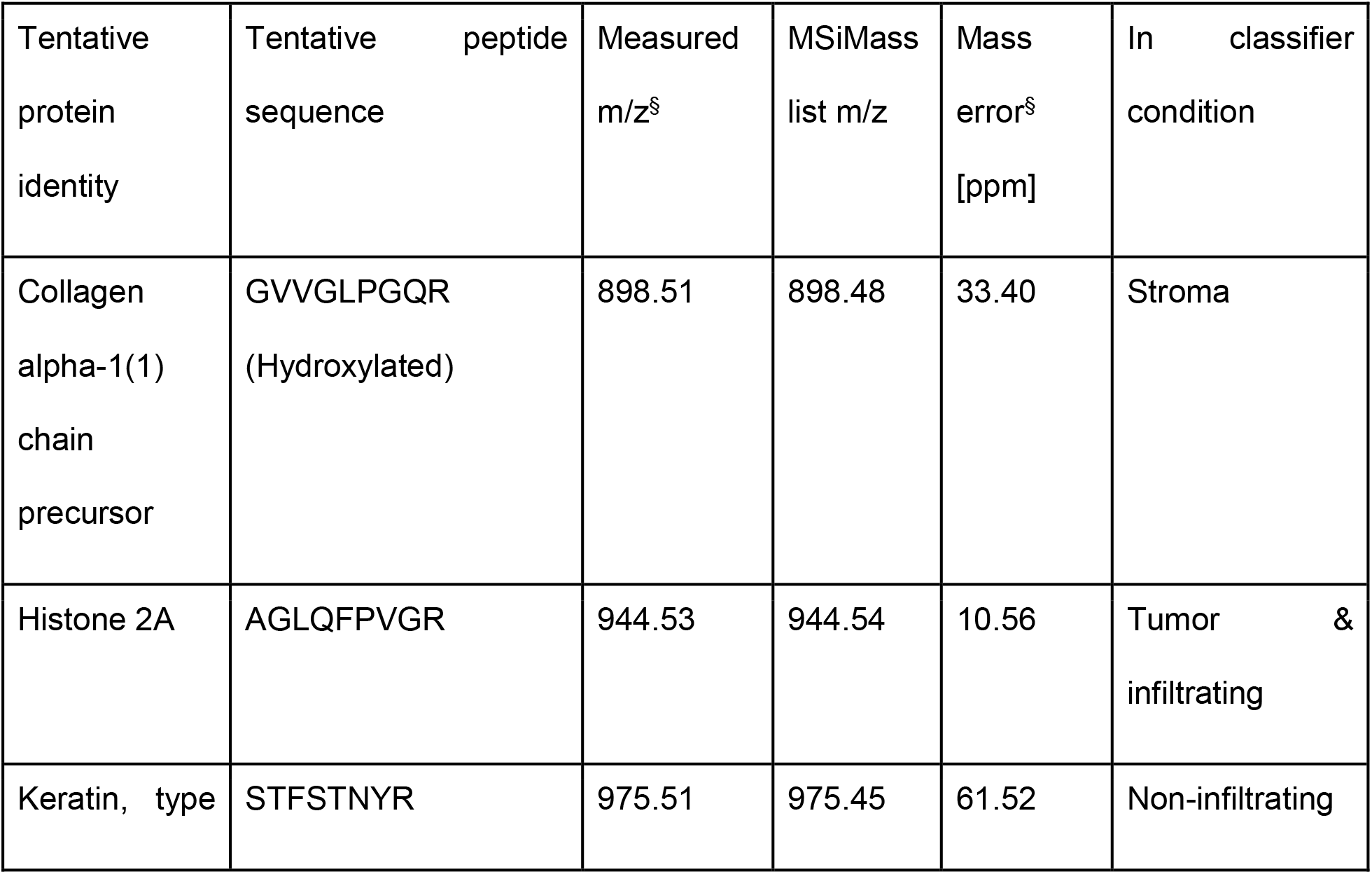
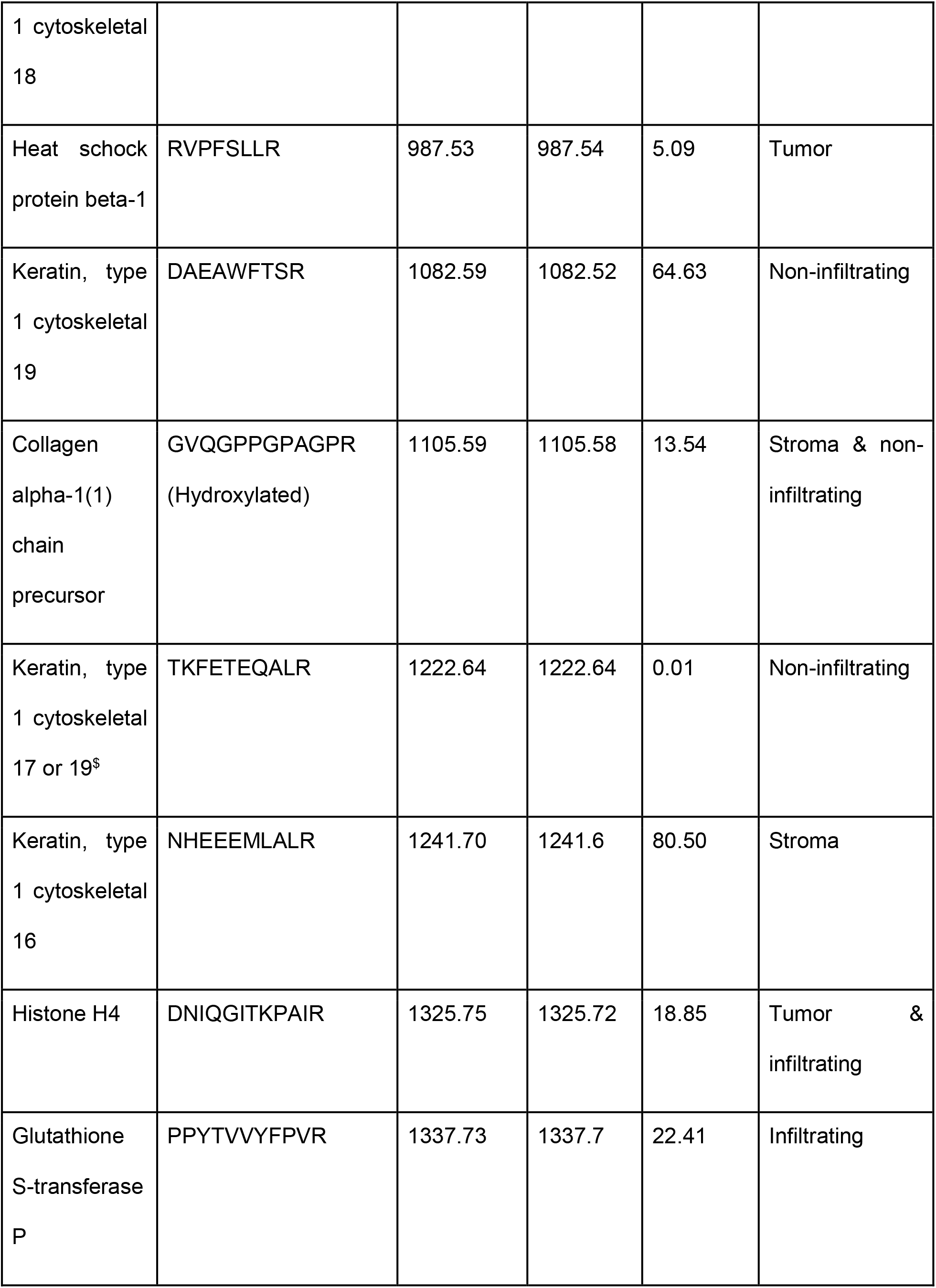
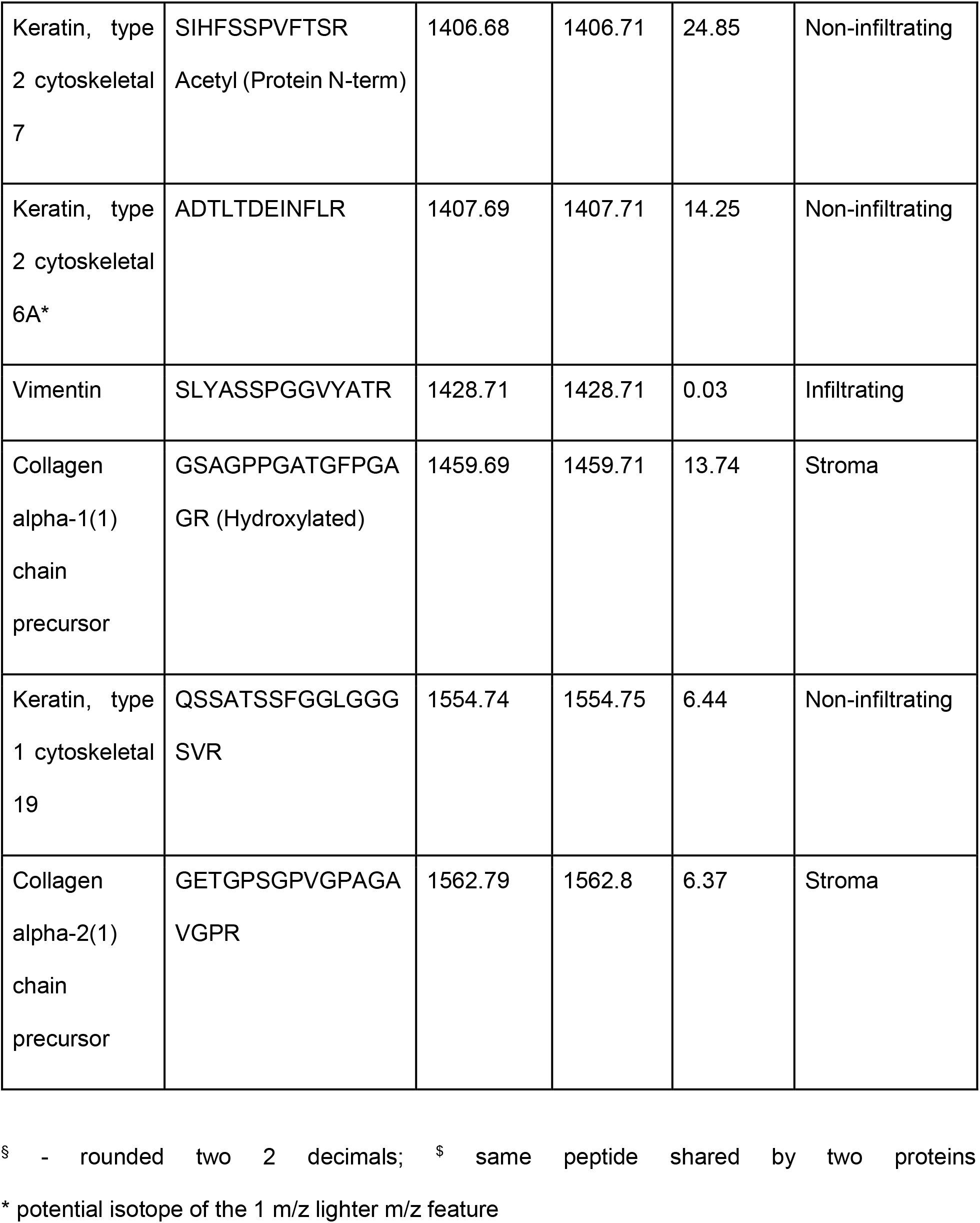
Peptide m/z matches obtained via the MSiMass list

The tentative m/z identifications of the tumor-stroma classifier point towards ubiquitous peptides that are likely to be found in other solid tumors and their surrounding stroma as well. Histone 2A and H4 were part of the tumor classification and likely indicators of increased cell density in the urothelium (transitional epithelium) compared to stroma tissue, because their abundance is proportional to the amount of DNA [34]. Another hit for the tumor classifier was heat shock protein beta-1, which is a member of the heat shock protein family, which has been linked to (urothelial) cancers before [35,36]. Stroma is connective tissue and therefore characterized by protein fibrils made for example out of collagens. This intrinsically corroborates collagen alpha-1(1) and alpha-2(1) chain precursors, which we found to be important for stromal classification. Cytokeratin 16 has been associated with ureter, bladder and urethra and also keratinization of urothelial carcinomas [37]. However, as an epithelial cell specific intermediate filament, it is probably a misidentification as it was part of the stromal classifier.

The obtained tentative identities for the muscle-infiltrating vs. non-muscle infiltrating classifier showed that several cytokeratins are associated with non-muscle invasive low-grade urothelial cancer, while histone 2A, histone H4, vimentin and glutathione S-transferase P were predictors for muscle-infiltrating urothelial carcinoma.

## DISCUSSION

We successfully conduct a fully transparent and reproducible analysis of a urothelial cancer MSI study in the Galaxy framework. The complete analysis was performed on a single platform, the European Galaxy Server. The previously established Galaxy MSI tools [18] allowed for all necessary analysis steps to classify different tissue types and tumor subtypes of an urothelial cancer cohort. This included co-registration of optical and MSI images, thorough quality controls, pre-processing as well as statistical modelling and peptide identification. Technical artifacts such as intensity batch effects and m/z shifts could be observed and removed via the ‘MSI qualitycontrol’ tool in combination with an adjusted pre-processing. Classification of different tissue types as well as different urothelial tumor subtypes was achieved with high accuracy and a few m/z features could be assigned to tentative identifications, which mainly were in line with the biological context.

All Galaxy analysis histories were published. They contain raw, meta and intermediate results data, as well as all tool names, tool versions, and all set tool parameters. Thus, every detail of the performed analysis can be re-traced and reproduced. In a copy of the Galaxy histories, researchers can adjust the analysis procedure according to their interest and inspect how changing different steps or parameters will change the outcome. Even though the Galaxy analysis history alone enables full reproducibility, we published all raw data including pathological annotations of stained tissues in the PRIDE proteomics data repository. This allows re-use of the data for new urothelial carcinoma studies and fosters future bioinformatic investigations since it represents the first human peptide imaging study that contains different disease groups and releases spectra wise pathological annotations for a complete patient cohort. While we have used the European Galaxy server for the analysis, studies with stricter data security restrictions could perform the analysis via ready to use docker containers on their local computing infrastructure [18]. However, to increase the trust in published MSI studies and to forward the MSI field it will become increasingly important to share raw data and analysis code, which requires to include data sharing into ethic approvals and patient consent forms from the beginning on.

## CONCLUSION

We have performed the complete analysis of an urothelial cancer cohort in a single platform, the European Galaxy server. Having used an outdated mass spectrometer, our study shows the importance of quality controls and pre-processing adjustment in order to detect and remove technical artifacts. Afterwards, we were able to classify tumor and stroma tissues as well as muscle-infiltrating and non-muscle invasive urothelial carcinomas based on their tryptic peptide composition with high accuracy and biologically explainable peptide identifications. In addition to the translational and biological findings, we highlight the potential for translational MSI studies and set new levels in terms of reproducibility and transparency by sharing all raw data and spectra annotations as well as the complete analysis histories. We would like to encourage the community to join our efforts to lay the foundation for advancing MSI towards clinical settings.

## Abbreviations

FAIR: findability, accessibility, interoperability, and reusability
MALDI: matrix assisted laser desorption/ionization
MS: mass spectrometry
MSI: mass spectrometry imaging
FFPE: formalin-fixed paraffin embedded SSC = spatial shrunken centroids
TMA: tissue microarrays
TOF: time of flight

## Declarations

### Ethics approval and consent to participate

The study was approved by the Ethics Committee of the University Medical Center Freiburg (no. 491/16) and all patients gave written informed consent.

### Consent for publication

Not applicable.

### Availability of data and materials

The mass spectrometry imaging raw data and stained optical images and annotated regions of interest are available in the ProteomeXchange Consortium via the PRIDE [30] partner repository with the dataset identifier PXD026459 (https://www.ebi.ac.uk/pride/archive/projects/PXD026459).

Galaxy workflows as well as all analysis histories are available via the European Galaxy server: https://github.com/foellmelanie/Bladder_MSI_Manuscript_Galaxy_links.

### Competing interests

The authors declare that they have no competing interests.

### Funding

OS acknowledges funding by the Deutsche Forschungsgemeinschaft (DFG, SCHI 871/11-1, SCHI 871/15-1, GR 4553/5-1, PA 2807/3-1, INST 39/1244-1 (P12), INST 39/766-3 (Z1), GRK 2606 “ProtPath”), the ERA PerMed programs (BMBF, 01KU1916, 01KU1915A), the German-Israel Foundation (grant no. 1444), and the German Consortium for Translational Cancer Research (project Im- pro-Rec). OV acknowledges funding by NSF-BIO/DBI (1950412) and NIH-NLM-R01 (1R01LM013115). PB acknowledges funding by the Fördergesellschaft Forschung Tumorbiologie and the Braun Stiftung.

### Authors’ contributions

PB and OS conceptualized and supervised the study. KW selected patients and established the study cohort. VV prepared the samples, carried out the mass spectrometry imaging measurements and the tissue staining. MCF analyzed the data. DG and OV contributed to statistical data analysis. KEA and PB annotated the pathological regions of interest. MCF, KEA, DG, OV, PB and OS interpreted the data. MCF prepared figures, tables and wrote the manuscript. All authors read and approved the final manuscript.

## Acknowledgements

The authors thank Lennart Moritz and Julia Huber for their excellent technical assistance. The authors acknowledge the support of the Freiburg Galaxy Team: Björn Grüning and Prof. Rolf Backofen, Bioinformatics, University of Freiburg, Germany funded by Collaborative Research Centre 992 Medical Epigenetics (DFG Grant SFB 992/1 2012) and German Federal Ministry of Education and Research (BMBF Grant 031 A538A RBC (de.NBI)).

## Notes

### Competing Interest Statement

The authors have declared no competing interest.

## References

1. Aichler M, Walch A. MALDI Imaging mass spectrometry: current frontiers and perspectives in pathology research and practice. Lab Investig. Nature Publishing Group; 2015;95:422–31.

2. Vaysse PM, Heeren RMA, Porta T, Balluff B. Mass spectrometry imaging for clinical research-latest developments, applications, and current limitations. Analyst. 2017;142:2690–712.

3. Arentz G, Mittal P, Zhang C, Ho Y-Y, Briggs M, Winderbaum L, et al. Applications of Mass Spectrometry Imaging to Cancer. Adv Cancer Res. 2017;134.

4. Berghmans E, Boonen K, Maes E, Mertens I, Pauwels P, Baggerman G. Implementation of Maldi mass spectrometry imaging in cancer proteomics research: Applications and challenges. J. Pers. Med. MDPI AG; 2020. p. 1–12.

5. Meding S, Nitsche U, Balluff B, Elsner M, Rauser S, Schöne C, et al. Tumor Classification of Six Common Cancer Types Based on Proteomic Profiling by MALDI Imaging. J Proteome Res. 2012;11:1996–2003.

6. Kriegsmann M, Casadonte R, Kriegsmann J, Dienemann H, Schirmacher P, Hendrik Kobarg J, et al. Reliable Entity Subtyping in Non-small Cell Lung Cancer by Matrix-assisted Laser Desorption/Ionization Imaging Mass Spectrometry on Formalin-fixed Paraffin-embedded Tissue Specimens. Mol Cell Proteomics. 2016;15:3081–9.

7. Möginger U, Marcussen N, Jensen ON. Histo-molecular differentiation of renal cancer subtypes by mass spectrometry imaging and rapid proteome profiling of formalin-fixed paraffin-embedded tumor tissue sections. Oncotarget. 2020;11:3998–4015.

8. Oppenheimer SR, Mi D, Sanders ME, Caprioli RM. Molecular analysis of tumor margins by MALDI mass spectrometry in renal carcinoma. J Proteome Res. 2010;9:2182–90.

9. Balluff B, Frese CK, Maier SK, Schöne C, Kuster B, Schmitt M, et al. De novo discovery of phenotypic intratumour heterogeneity using imaging mass spectrometry. J Pathol. 2015;235:3–13.

10. Mittal P, Klingler-Hoffmann M, Arentz G, Winderbaum L, Lokman NA, Zhang C, et al. Lymph node metastasis of primary endometrial cancers: Associated proteins revealed by MALDI imaging. Proteomics. 2016;16:1793–801.

11. Hoffmann F, Umbreit C, Krüger T, Pelzel D, Ernst G, Kniemeyer O, et al. Identification of Proteomic Markers in Head and Neck Cancer Using MALDI–MS Imaging, LC–MS/MS, and Immunohistochemistry. Proteomics - Clin Appl. 2019;13:1–10.

12. Cazares LH, Troyer D, Mendrinos S, Lance RA, Nyalwidhe JO, Beydoun HA, et al. Imaging mass spectrometry of a specific fragment of mitogen-activated protein kinase/extracellular signal-regulated kinase kinase kinase 2 discriminates cancer from uninvolved prostate tissue. Clin Cancer Res. 2009;15:5541–51.

13. Erich K, Sammour DA, Marx A, Hopf C. Scores for standardization of on-tissue digestion of formalin-fixed paraffin-embedded tissue in MALDI-MS imaging. Biochim Biophys Acta - Proteins Proteomics. Elsevier B.V1.; 2017;1865:907–15.

14. Ly A, Longuespée R, Casadonte R, Wandernoth P, Schwamborn K, Bollwein C, et al. Site- to- Site Reproducibility and Spatial Resolution in MALDI–MSI of Peptides from Formalin- Fixed Paraffin- Embedded Samples. PROTEOMICS – Clin Appl. 2019;13:1800029.

15. Buck A, Heijs B, Beine B, Schepers J, Cassese A, Heeren RMA, et al. Round robin study of formalin-fixed paraffin-embedded tissues in mass spectrometry imaging. Analytical and Bioanalytical Chemistry; 2018;5969–80.

16. Gustafsson OJR, Winderbaum LJ, Condina MR, Boughton BA, Hamilton BR, Undheim EAB, et al. Balancing sufficiency and impact in reporting standards for mass spectrometry imaging experiments. Gigascience. 2018;7:1–13.

17. Gustafsson JOR, Eddes JS, Meding S, Koudelka T, Oehler MK, McColl SR, et al. Internal calibrants allow high accuracy peptide matching between MALDI imaging MS and LC-MS/MS. J Proteomics. Elsevier B.V.; 2012;75:5093–105.

18. Öll MC, Moritz L, Wollmann T, Stillger MN, Vockert N, Werner M, et al. Accessible and reproducible mass spectrometry imaging data analysis in Galaxy. Gigascience. 2019;8:628719.

19. Afgan E, Baker D, Batut B, van den Beek M, Bouvier D, Čech M, et al. The Galaxy platform for accessible, reproducible and collaborative biomedical analyses: 2018 update. Nucleic Acids Res. Oxford University Press; 2018;46:W537–44.

20. Wilkinson MD. Comment: The fair guiding principles for scientific data management and stewardship. Sci Data. 2016;1–9.

21. Bronsert P, Weißer J, Biniossek ML, Kuehs M, Mayer B, Drendel V, et al. Impact of routinely employed procedures for tissue processing on the proteomic analysis of formalin-fixed paraffin-embedded tissue. Proteomics - Clin Appl. 2014;8:796–804.

22. Gustafsson JOR, Oehler MK, McColl SR, Hoffmann P. Citric acid antigen retrieval (CAAR) for tryptic peptide imaging directly on archived formalin-fixed paraffin-embedded tissue. J Proteome Res. 2010;9:4315–28.

23. Stoeckli M, Staab D, Wetzel M, Brechbuehl M. iMatrixSpray: A Free and Open Source Sample Preparation Device for Mass Spectrometric Imaging. Chim Int J Chem. 2014;68:146–9.

24. European Galaxy Instance. [cited 2019 Mar 9];

25. Bemis KD, Harry A, Eberlin LS, Ferreira C, Van De Ven SM, Mallick P, et al. Cardinal: An R package for statistical analysis of mass spectrometry-based imaging experiments. Bioinformatics. 2015;31:2418–20.

26. Keller BO, Sui J, Young AB, Whittal RM. Interferences and contaminants encountered in modern mass spectrometry. Anal Chim Acta. 2008;627:71–81.

27. Bemis KD, Harry A, Eberlin LS, Ferreira CR, van de Ven SM, Mallick P, et al. Probabilistic Segmentation of Mass Spectrometry (MS) Images Helps Select Important Ions and Characterize Confidence in the Resulting Segments. Mol Cell Proteomics. 2016;15:1761–72.

28. Pedregosa F, Varoquaux G, Gramfort A, Michel V, Thirion B, Grisel O, et al. Scikit-learn: Machine Learning in Python. J Mach Learn Res. 2011;12:2825–30.

29. McDonnell LA, Walch A, Stoeckli M, Corthals GL. MSiMass list: A public database of identifications for protein MALDI MS imaging. J Proteome Res. 2014;13:1138–42.

30. Vizcaíno JA, Csordas A, Del-Toro N, Dianes JA, Griss J, Lavidas I, et al. 2016 update of the PRIDE database and its related tools. Nucleic Acids Res. 2016;44:D447–56.

31. Deininger S-O, Cornett DS, Paape R, Becker M, Pineau C, Rauser S, et al. Normalization in MALDI-TOF imaging datasets of proteins: practical considerations. Anal Bioanal Chem. 2011;401:167–81.

32. Fonville JM, Carter C, Cloarec O, Nicholson JK, Lindon JC, Bunch J, et al. Robust data processing and normalization strategy for MALDI mass spectrometric imaging. Anal Chem. 2012;84:1310–9.

33. Lopez-Beltran A, Scarpelli M, Montironi R, Kirkali Z. 2004 WHO Classification of the Renal Tumors of the Adults. Eur Urol. 2006;49:798–805.

34. Wiśniewski JR, Hein MY, Cox J, Mann M. A “proteomic ruler” for protein copy number and concentration estimation without spike-in standards. Mol Cell Proteomics. 2014;3497–506.

35. Chatterjee S, Burns T. Targeting Heat Shock Proteins in Cancer: A Promising Therapeutic Approach. Int J Mol Sci. 2017;18:1978.

36. Ischia J, So AI. The role of heat shock proteins in bladder cancer. Nat Rev Urol. 2013;10:386–95.

37. Bernot KM, Coulombe PA, McGowan KM. Keratin 16 Expression Defines a Subset of Epithelial Cells During Skin Morphogenesis and the Hair Cycle. J Invest Dermatol. 2002;119:1137–49.

